# Sirt3 mediated by lentiviral vector in the periaqueductal gray suppresses morphine withdrawal in rats: A preliminary study

**DOI:** 10.1101/743062

**Authors:** Shue Liu, Hyun Yi, Jun Gu, Daigo Ikegami, Kentaro Hayashi, Shuanglin Hao

## Abstract

Opioid use disorder (OUD) is a significant clinical and social problem, inducing dependence/addiction and over-dose death. Opioid dependence/withdrawal contributes to the addiction vulnerability. Limited understanding of the exact mechanisms of morphine withdrawal leads to failure to adequately manage opioid withdrawal symptoms. Determining new molecular mechanisms of morphine withdrawal (MW) may allow development of novel therapeutic strategies for treating this disorder. Chronic morphine with naloxone precipitation induces MW behavioral response. Sirt3 (one member of sirtuins family) as a mitochondrial fidelity, plays an important role in mitochondrial homeostasis through the direct regulation of mitochondrial energy metabolism, ATP synthesis, detoxification of mitochondrial ROS, etc. In the pilot study, we found that (1) cultured neurons infected with lentiviral vector expressing Sirt3 induced over-expression of Sirt3, (2) microinjection of LV-Sirt3 into the vlPAG increased Sirt3 protein expression in rats, (3) MW lowered the expression of Sirt3 in the vlPAG, and (4) microinjection of LV-Sirt3 into the vlPAG decreased the MW behavioral response. Current preliminary study demonstrates that complement of Sirt3 in the PAG suppressed MW, providing a novel therapeutic approach to morphine physical withdrawal symptoms. The exact up-and/or down-stream factors of Sirt3 in the model are under the investigation.

## 1. Introduction

Opioid use disorders (OUD) have reached an epidemic level in the United States ^1^. Opioid use disorder (OUD) is a significant clinical and social problem, inducing dependence/addiction and over-dose death^2^. CDC Morbidity and Mortality Weekly Report shows that from 2016 to 2017, synthetic opioid-involved overdose death rates increased 45.2% ^3^. Repeated opioid use induces hyperalgesia, tolerance, dependence and addiction ^4–8^. There is an increased risk of overdose death with higher daily opioid doses for increasing analgesic effect ^9^. Opioid physical dependence is one of the determinants of opiate abuse in dependent individuals. Physical dependence to opiates can be demonstrated by either termination of drug administration or opiate antagonist precipitation of the withdrawal syndrome. Existing pharmacotherapies for opioid dependence management may have high rates of failure (e.g. relapse and/or dropout)^10, 11^ because the exact mechanisms of opioid dependence are still poorly understood.

Neuropharmacological studies show that several regions (e.g. locus ceruleus (LC), periaqueductal gray (PAG)) of the brain with high opioid receptors, are related to MW or drug abuse ^12–14^. Functional and anatomical studies have implicated an important role for the PAG in opioid withdrawal ^15–18^. Sirtuins are protein deacetylases dependent on nicotine adenine dinucleotide (NAD) in organisms ranging from bacteria to humans^19^. In mitochondria, some specific fidelity proteins activate signaling pathways to maintain mitochondrial homeostatic poise. Sirtuin 3 (Sirt3, one of sirtuins family), a major deacetylase in mitochondria, is essential in maintaining mitochondria homeostasis. Sirt3 is localized into the inner mitochondrial membrane. Sirt3, a fidelity protein for mitochondrial function activity ^20^, has recently received great attention, however, its role in the PAG in MW is still not clear.

In the preliminary study, we used lentiviral vector-mediated expression of human Sirt3 to determine the effect of Sirt3 overexpression in the ventrolateral PAG (vlPAG) on naloxone-precipitated MW in rats. In the near future, we will determine the exact molecular mechanisms of up-and down-stream pathway of Sirt3 in the model.

## 2. Materials and methods

### Construction of the lentiviral vector expressing Sirt3

We constructed a lentiviral vector, human Sirt3 expressing lentiviral vector with lentiviral plasmid, packaging plasmid, and VSV-G envelope generated by inserting human Sirt3 plasmid (Purchased from Addgene #13814) into HIV.SIN.cPPT.CMV.eGFP.WPRE (Purchased from U Penn Vector Core, Philadelphia, PA) at the EcoIR site (LV-Sirt3, packaged by Dr. Mingjie Li, Neurology, Washington University School of Medicine, St. Louis, MO). Control lentivirus (LV-GFP) was made but no human sirt3 gene.

### Animal and evaluation of chronic morphine physical dependence

Male 7-8 weeks Sprague-Dawley rats (body weight 225-250 g) were housed two per cage approximately 1 week prior to the beginning of the research, with free access to food and water and maintained on a 12:12, light: dark schedule at 21 °C and 60% humidity. All housing conditions and experimental procedures were approved by the University Animal Care and Use Committee. Morphine withdrawal procedure was induced as described previously ^21^. Rats received escalating doses of intraperitoneal (IP) morphine for a period of 5 days as follows: day 1, 10 mg/kg (8AM, IP) and 15 mg/kg (8PM); day 2, 20 and 25 mg/kg; day 3, 30 and 35 mg/kg; day 4, 40 and 45 mg/kg. On day 5, animals received a morning injection of 50 mg/kg, and 1 hour later, naloxone (4 mg/kg, IP) was administered to precipitate a MW response. Rats were placed individually in a test chamber (50×35×45 cm) immediately after naloxone, and we evaluated MW behavioral signs over the course of 30 min. Two types of signs were measured during abstinence, as described previously ^22, 23^. Episodes of wet-dog shakes and jumps were counted (i.e. recorded quantitatively); teeth chatter (vacuous chewing), diarrhea, rhinorrhea, ptosis, lacrimation, escaping, penile erection, and abnormal posture were evaluated over 5 min periods with one point being given for the presence of each sign during each period. We recorded the body weight of each rat before the injection of naloxone and then again at 60 min after naloxone. The scored signs defined in our previous work^24^, were based on our previous experience as modified from those described by Punch et al ^13^. A global withdrawal score was calculated for each rat by assigning a combination of the various physical signs of withdrawal ^25^. MW behavior counting were carried out by a blinded operator.

### Microinjection of lentiviral vector into vlPAG

For intracranial vector administration, anesthetized rats with isoflurane were placed in a stereotaxic headholder (David Kopf Instruments, Tujunga, CA). After the skull was exposed, a 33-gauge needle with microsyringe was directed bilaterally toward the vlPAG (AP −8.2 mm using bregma as 0, ML ±0.75 mm, DV −6.2 mm from the base of the skull). Rats were randomly divided to receive a bilateral vlPAG injection of LV vector expressing Sirt3 (LV-Sirt3, 1 µl of LV-GFP (3.4×10^8^ IU/ml)) or control vector expressing GFP (LV-GFP) with a micropump (0.5µl/min). After injection, the injector was left in situ for an additional 4 min before it was removed. Animals showing neurological deficits or 85 % loss of body weight after microinjection were excluded.

### Western blot

Rat brains were harvested under deep anesthesia. A tissue block including a segment at the level of the vlPAG was cut on an ice cold glass plate ^16^. The vlPAG from the tissue block was harvested by taking punches with a 14 gauge puncture needle as described previously ^26^. The punched vlPAG tissue was homogenized with 100 µl of ice-cold lysis buffer (150 mM sodium chloride, 1.0% NP-40, 0.5% sodium deoxycholate, 0.1% SDS, 50 mM Tris, pH 8.0) containing protease inhibitors (Sigma, St Louis, MO) and phosphatase inhibitor cocktail 2 and 3 (Sigma, St Louis, MO). We sonicated tissue for homogenates and then centrifuged it at 15,000 g for 20 min at 4 °C. The supernatant was collected and assayed for protein content using the BCA assay method (Pierce, Rockford, IL, USA) and stored at −80 °C until further use. Protein (∼20 µg) was electrophoresed on a 12% SDS-PAGE gel, then transferred to a PVDF membrane, and blocked with 1x Rapid block solution (Amresco, Fountain Parkway Solon, OH) at room temperature for one hour. The primary antibodies, rabbit anti-Sirt3(cata#2627, 1:2000, Cell signaling, Danver, MA) and mouse β-actin(cata#A5441, 1:8000, Sigma, St. Louis, MO) were incubated overnight at 4 °C in fresh blocking buffer. The membrane was washed and incubated with complementary secondary antibodies (1:4000, horseradish peroxidase conjugated IgG antibody, Santa Cruz biotechnology, Dallas, TX) for 1 h at room temperature. The membranes were washed in washing buffer for 10 minutes three times and the antibodies were then revealed using Super Signal west dura extended duration substrate (Thermo Fisher Scientific Inc, Rockford, IL, USA). For densitometric analyses, blots were quantified with Quantity One analysis software (Bio-Rad, Hercules, CA). The results were expressed as the ratio to β-actin immunoreactivity.

### Immunohistochemistry

For the PAG immunohistochemistry, rats were perfused intracardially with 4% PFA in 0.1 M phosphate buffer, and the brains were removed, postfixed in the same solution for overnight, and cryoprotected with 30% sucrose in phosphate-buffered solution for 2 days. In order to define the co-localization with NeuN, Sirt3, GFAP, or Iba1, cryostat sections (25 μm thickness) were incubated 48 h at 4 °C with primary antibodies, mouse anti-NeuN (cata #MAB377, 1:500, Millipore, Billerica, MA), rabbit anti-Sirt3(cata#2627, 1:100, Cell signaling, Danvers, MA), mouse anti-GFAP (cata. # G3893, 1: 3000, Sigma, St Louis, MO), and rabbit anti-Iba1 (cata.# 019-19741, 1:2000, Wako, Richmond, VA) followed by fluorescent IgG with Alexa Fluor 488 or 594 (1:1000, Invitrogen Life Technologies) for 2 hours at room temperature. Fluorescence images were captured by a fluorescent microscopy (Fluorescent M Leica/Micro CDMI 6000B).

### Sirt3 expression in cultured neuronal cells treated with lentiviral vectors

The rat neuronal cell line (B35, CRL-2754) was obtained from American Type Culture Collection (ATCC, Manassas, VA). For western blots, the cells were plated in a 6-well plate, infected with either LV-Sirt3 or LV-GFP at a multiplicity of infection (MOI, =1) for 3 days, and collected to measure human Sirt3 expression using western blot.

### Drugs and data analysis

Morphine sulfate was obtained from West-Ward Pharmaceuticals, Eatontown, NJ. Naloxone hydrochloride was obtained from Sigma. Naloxone was injected intraperitoneally in a volume of 1 ml/kg of body weight. All drugs were dissolved in physiologic (0.9%) saline. The sample size estimate was based on our previous studies^16, 27^. The statistical significance of the differences was determined by one-way ANOVA (SPSS21, IBM, Armonk, NY, or StatView5, SAS Institute Inc. Cary, NC), or *t* test. The difference between the time-course curves of MW behavior was determined using 2-way ANOVA repeated measure (GraphPad Prism 5). *P*-values of less than 0.05 were considered to be statistically significant.

## 3. Results

### 3.1. Sirt3 expression mediated by lentiviral vectors *in vitro* and *in vivo*

Three days after transfection with LV-Sirt3 or control vector LV-GFP, neurons were collected, and Sirt3 were detected by western blot. We found that Sirt3 expression in neurons infected with LV-Sirt3 was significantly higher compared to LV-GFP (Figure 1).

**Figure 1.**
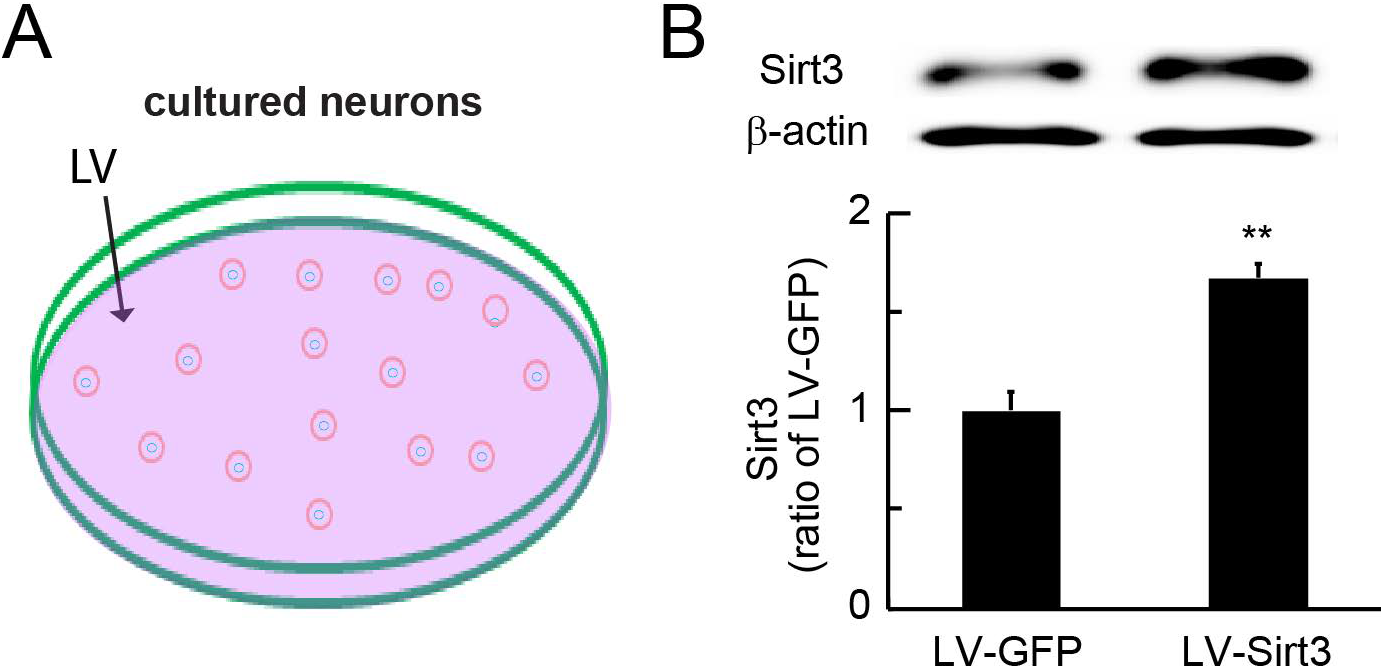
Sirt3 expression mediated by lentiviral vectors *in vitro*. Three days after transduction with LV-Sirt3 or control vector LV-GFP (**A**), neurons were collected, and Sirt3 was detected by western blot (**B**). Sirt3 expression in neurons infected with LV-Sirt3 was significantly higher compared to LV-GFP, \*\**P*<0.01, *t* test, n=6.

In *in vivo* study, Figure 2A showed that the track of vlPAG injection was cleanly seen at the vlPAG, marked with red ellipse. To evaluate the expression of Sirt3 mediated by LV-Sirt3, 2 weeks after microinjection of LV-Sirt3 into the vlPAG, the vlPAG tissue was collected, and Sirt3 expression was compared using western blot. Western blot showed that Sirt3 expression in LV-Sirt3-microinjected animals was higher compared to LV-GFP (Figure 2B).

**Figure 2.**
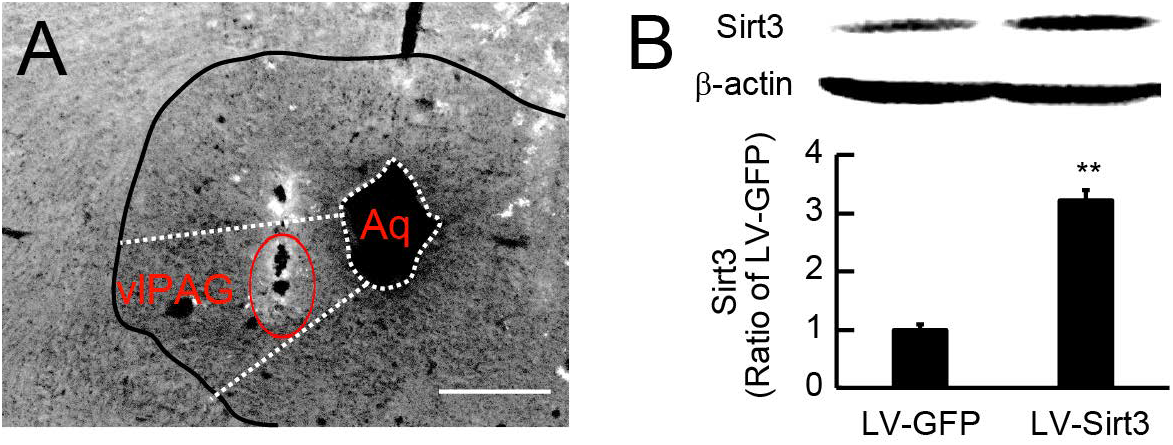
Sirt3 expression mediated by lentiviral vectors *in vivo*. (**A**) The track of vlPAG injection was cleanly seen at the vlPAG, marked with red ellipse. (**B**) Western blot showed that substantial amounts of Sirt3 in LV-Sirt3-microinjected animals compared to LV-GFP, 2 week after microinjection of LV-Sirt3 into the vlPAG, \*\**P*<0.01, *t* test, n=4-5.

### 3.2. MW lowered the expression of Sirt3 at the vlPAG

We have reported that chronic escalating morphine with naloxone precipitation induced morphine withdrawal response ^16^. It is suggested that oxidative stress is involved in the development of morphine physical dependence. Heroin decreased the total antioxidant capacity of serum and the anti-oxidative enzymes activities in brains ^28^. Sirt3 can rescue neuronal loss in various neurodegenerative models^29^. Loss of Sirt3 increases mtO_2_^•−^ levels, whereas overexpression of Sirt3 decreases mtO_2_^•−^ levels ^30, 31^. Our unpublished data showed that MW increased mitochondrial superoxide in the vlPAG. Here, in chronic morphine with naloxone precipitation model, we observed that Sirt3-immunoreactivity expression mainly was co-localized with NeuN (a marker of neurons) (Figure 3B-D), but not GFAP (a marker of astrocytes) or Iba1 (a marker of microglia) (*data not shown*). Western blots demonstrated that the expression of Sirt3 in MW group was lower than that in the sham group (Figure 3E).

**Figure 3.**
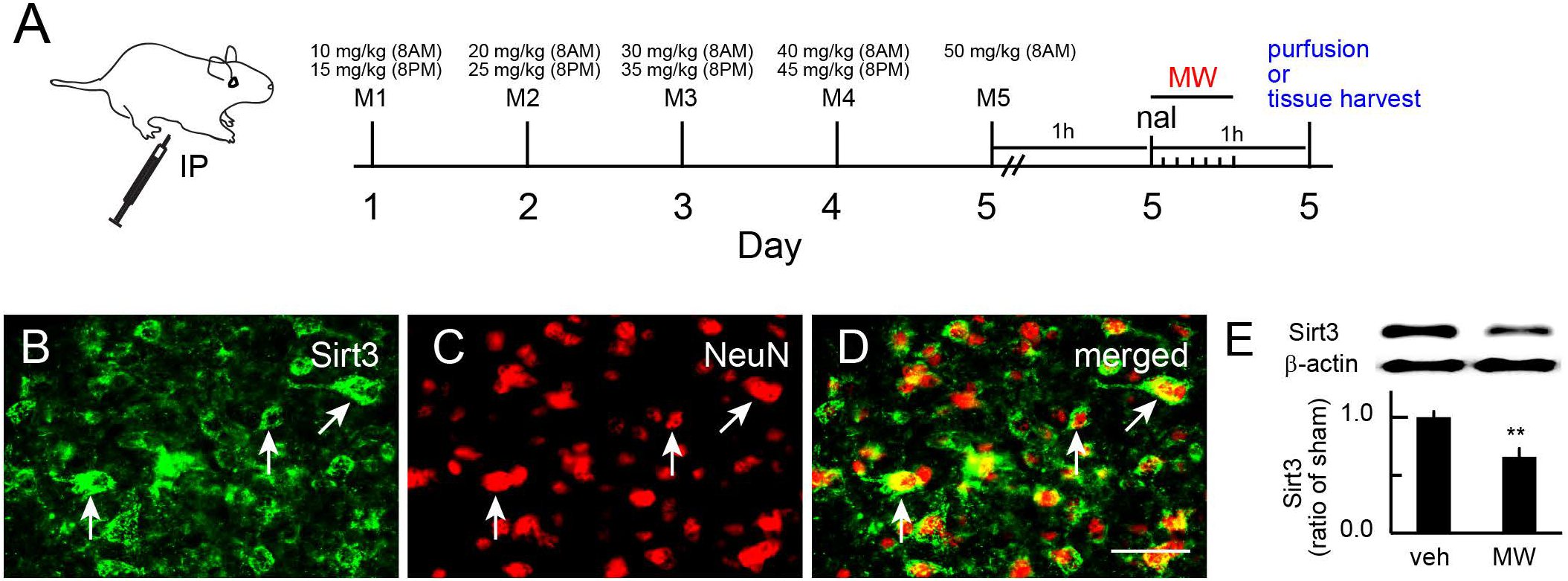
MW lowered Sirt3 expression in the vlPAG. (**A**) The MW procedure was shown. (**B-D**) Double immunostaining showed that Sirt3-immunoreactivity mainly was co-localized with NeuN (a marker of neurons). (E) Western blots demonstrated that the expression of Sirt3 in MW group was lower than that in the sham group, \*\**P*<0.01, *t* test, n=6 (Figure 3D). nal, naloxone. Scale bar, 50µm.

### 3.3. Neuronal-targeting GFP expression mediated by lentiviral vectors microinjected into the rat PAG

We constructed the lentiviral vector to express human *sirt3* gene (LV-Sirt3). Control vector contains gfp gene but no human Sirt3 gene (LV-GFP). We microinjected LV-GFP into the PAG. Ten days after injection, rats were perfused with 4% PFA. The direct green fluorescence was detected, and GFP was co-localized with NeuN immunostaining (Figure 4), but not glia markers GFAP or Iba1 (Figure 4).

**Figure 4.**
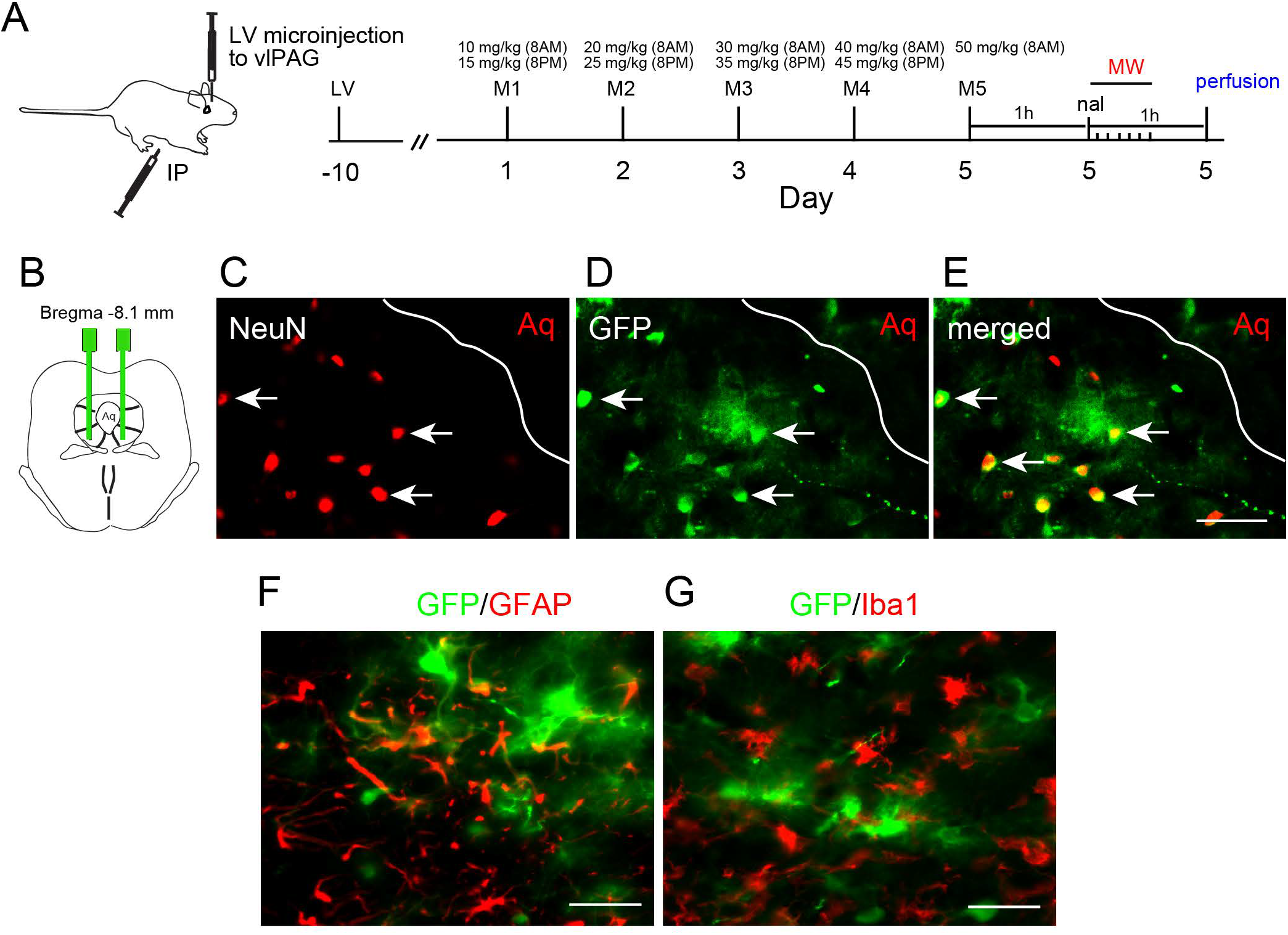
Neuronal-targeting GFP expression mediated by lentiviral vectors microinjected into the rat PAG. (**A**) The LV microinjection and the MW procedure was shown. (**B-E**) We microinjected LV-GFP into the PAG. Ten days after injection, rats were perfused with 4% PFA. The direct green fluorescence was detected, and GFP was co-localized with NeuN immunostaining, but not glia markers GFAP (**F**) or Iba1 (**G**). LV, lentiviral vector; IP, intraperitoneal injection. Scale bar, 50µm.

### 3.4. Lentiviral vector-mediated expression of Sirt3 suppressed MW behavioral response

LV-SIRT3 or LV-GFP was bilaterally microinjected into the vlPAG. For MW induction, ten days after vector microinjection, chronic escalating morphine or vehicle (sham) was administered. At day 5 of chronic morphine, one hour after the last injection of morphine, animals received naloxone. Withdrawal scores were calculated for each rat immediately after naloxone. In sham groups with microinjection of LV-Sirt3 or LV-GFP, there was no significant difference in the MW behavior scores (Figure 5).

**Figure 5.**
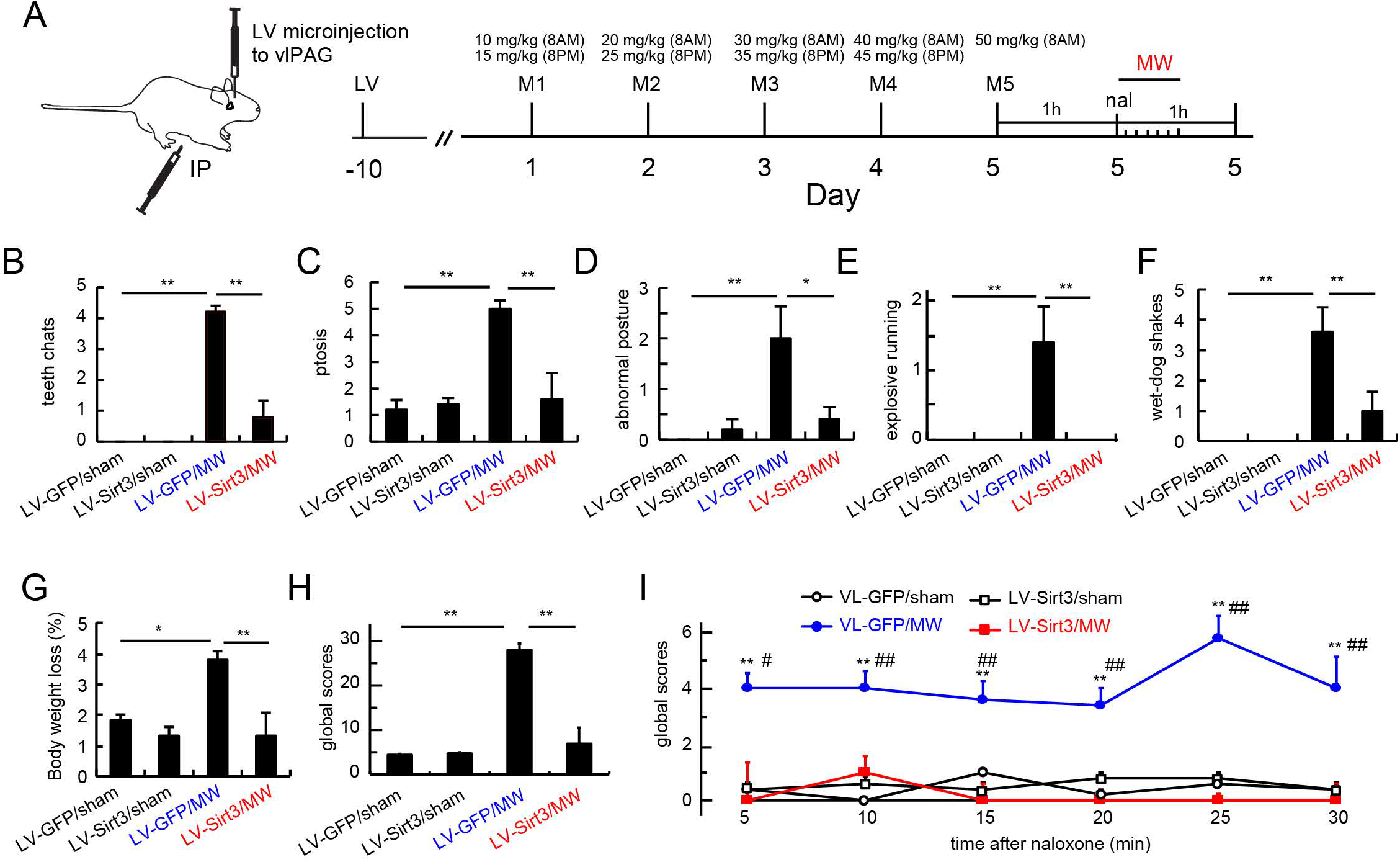
Lentiviral vector-mediated expression of Sirt3 suppressed MW behavioral response. (**A**) The LV microinjection and MW procedure was shown. Lentiviral vector LV-SIRT3 or LV-GFP was bilaterally microinjected into the vlPAG. For MW induction, ten days after vector microinjection, chronic escalating morphine or vehicle (sham) was administered. At day 5 of chronic morphine, one hour after the last injection of morphine, animals received naloxone. There was a significant increase in MW behavioral scores of teeth chat (**B**), ptosis (**C**), abnormal posture (**D**), explore running (**E**), wet-dog shakes (**F**), body weight loss (**G**), and global scores (**H**) in rats with LV-GFP/MW compared to LV-GFP/sham rats. MW behavioral scores of teeth chat, ptosis, abnormal posture, explore running, wet-dog shakes, body weight loss, and global scores in the LV-Sirt3/MW group, were significantly lower than that in the LV-GFP/MW group, one-way ANOWA Bonferroni test. (**I**) The time course of MW behavioral response, \*\**P*<0.01 vs. LV-GFP/sham, #*P*<0.05, ##*P*<0.01 vs. LV-Sirt3/MW, 2-way ANOVA Bonferroni test, n=5.

There was a significant increase in behavioral scores of teeth chat (Figure 5B), ptosis (Figure 5C), abnormal posture (Figure 5D), explore running (Figure 5E), wet-dog shakes (Figure 5F), body weight loss (Figure 5G), and global scores (Figure 5H) in rats with LV-GFP/MW compared to LV-GFP/sham rats. MW behavioral scores of teeth chat, ptosis, abnormal posture, explore running, wet-dog shakes, body weight loss, and global scores in the LV-Sirt3/MW group, were significantly lower than that in the LV-GFP/MW group (Figure 5B-H). For the comparison of the time course of MW behavioral response, 2-way ANOVA Bonferroni test showed that there was a significant increase in MW behavioral global scores in the LV-GFP/MW rats compared to the LV-GFP/sham group at time points of 5-30 min (P < 0.01) (Figure 5I), and that MW global scores in the LV-Sirt3/MW group were significantly lower than that in the LV-GFP/MW group (Figure 5I).

### 4. Discussion

OUD is a significant clinical and social problem, inducing dependence and over-dose death^2^. Elucidation of the exact mechanisms of MW is important to treatment of opioid dependence. In the pilot study, we found that (1) cultured neurons infected with lentiviral vector expressing Sirt3 induced over-expression of Sirt3, (2) microinjection of LV-Sirt3 into the vlPAG increased Sirt3 protein expression in rats, (3) MW lowered the expression of Sirt3 in the vlPAG, and (4) microinjection of LV-Sirt3 into the vlPAG decreased the MW behavioral response.

Neuropharmacological studies show that multiple regions of the brain with high opioid receptors, are related to MW or drug abuse in human and animals ^13, 15, 32–41^. Anatomical, and neurochemical studies have implicated an important role of the PAG in opioid withdrawal ^15–18, 42–44^. Distinct PAG columns project to midline and intralaminar thalamic regions ^45^. The lateral hypothalamic area projects selectively to the vlPAG ^46^. PAG projecting to several regions of the brain including the locus ceruleus (LC), may function as an integrating center to coordinate responses to opiate withdrawal ^47^.

Sirtuins are NAD^+^-dependent class III histone deacetylases, and they are present from bacteria to humans^48^. The mammalian sirtuins are related to wide activities including cellular stress resistance, genomic stability, gene silencing, tumorigenesis, and energy metabolism ^49^. Sirtuins are involved in cellular and nutrient stress inducing the activation of their deacetylase or ribosyltransferase activity^48, 50^. The sirtuins primarily target cellular proteins to regulate cell signaling networks ^49^ and post-translationally alter the activity of downstream protein targets^51^. Sirtuin activating compounds slow metazoan ageing by mechanisms that may be related to caloric restriction ^52^. Sirtuins modulate p53-mediated growth regulation^53^. Mammalian sirtuins contain 7 sirtuins (Sirt1-7) localized in the nucleus (Sirt1, 6, 7), cytoplasm (Sirt2), and mitochondria (Sirt3-5) ^54^. Sirt3 is the primary mitochondrial sirtuins. Calorie restriction reduces oxidative stress by Sirt3-mediated MnSOD activation ^55^. SIRT3(−/−) mouse embryonic fibroblasts (MEFs) exhibit abnormal mitochondrial physiology as well as increases in stress-induced superoxide levels and genomic instability, suggesting that SIRT3 is a genomically expressed, mitochondria-localized tumor suppressor ^56^. Mitochondria from Sirt3(−/−) animals display a selective inhibition of mitochondrial respiratory chain complex I activity; incubation of exogenous Sirt3 with mitochondria can augment Complex I activity, implicating that Sirt3 functions *in vivo* to regulate and maintain basal ATP levels^57^. Sirt3 as a mitochondrial fidelity maintains mitochondrial homeostasis through the direct regulation of mitochondrial energy metabolism, ATP synthesis, detoxification of mitochondrial ROS, etc.

Sirt3 is expressed at high levels in the central nervous system ^29^ and regulates every major aspect of mitochondrial biology, including reactive oxygen species (ROS) detoxification and mitochondrial dynamics ^58–60^. Superoxide radical (O_2_^•−^) can react with DNA, lipids, etc., to ultimately result in cell injury ^61^. Mitochondria are the major source of intracellular ROS in neurons ^62^. Sirt3 deacetylates MnSOD and thereby activates it for scavenging mtO_2_^•-30, 63–65^. Loss of Sirt3 increases mtO_2_^•−^ levels, whereas overexpression of Sirt3 decreases mtO_2_^•−^ levels^30, 31^. Sirt3 can rescue neuronal loss in various neurodegenerative models ^29^. Our current study demonstrates that MW decreased expression of Sirt3 in the PAG.

Oxidative stress is involved in the development of morphine physical dependence. Heroin decreased the total antioxidant capacity of serum and the anti-oxidative enzymes activities in brains ^28^. Pretreatment with antioxidants could inhibit oxidative stress, and alleviate opioid withdrawal in animals ^66–68^. Our unpublished data shows that MW increased the mitochondrial superoxide in the vlPAG and super-oxidative scavenger reduced MW behavioral response. We reported that herpes simplex viral vector over-expressing MnSOD in the vlPAG reduced MW behavioral response ^69^. It is highly possible that Sirt3 mediated by the lentiviral vector suppressed MW behavioral response through activating MnSOD.

### Future study

The pilot studies show that MW lowers neuronal Sirt3 expression in the PAG, suggesting that loss of sirt3 play an important role in the MW, however it is still not clear about the upstream pathway of Sirt3 loss. Our unpublished data showed that Sirt3 was mediated by EZH2 or miRNA in an epigenetic manner in chromic morphine or neuropathic pain state. Recent report shows that SUMO-specific protease SENP1 is accumulated in mitochondria and quickly de-SUMOylates and activates Sirt3 ^70^. Sirt3 also mediates the function/activity of MnSOD ^55, 71^. In the near future, we will determine the exact mechanisms of Sirt3 up-stream and down-stream factors in chromic morphine or neuropathic pain state.

## Acknowledgements

Supported by grants from the US National Institutes of Health R01NS066792 (S.H.), R01DA34749 (S.H.), R01DA047089 (S.H.), R01DA047157 (S.H.),

